# *EctoShed*: A novel *Gene Delivery Platform for Functional Analysis of Adipocyte-Shed Proteoforms*

**DOI:** 10.1101/2025.10.14.682314

**Authors:** Ana Rita Tavanez, Nadia Meincke Egedal, Natasa Stanic, Hande Topel, Jan-Wilhelm Kornfeld

**Author notes:** equal contribution (Ana Rita Tavanez, Nadia Meincke Egedal).

## Abstract

**Objective:** The proteolytic cleavage of membrane-bound proteins, ectodomain shedding, functionally expands the reservoire of proteins/peptides available for endocrine crosstalk, and metabolic regulation. However, the functional understanding of secreted proteoforms, including whether they act synergistically or antagonistically with their membrane precursors, is often unknown. We aimed to develop a novel viral vector based gene delivery platform enabling characterization of both membrane-bound and soluble proteoforms in adipocytes, independent of endogenous shedding.

**Results:** We describe a novel platform, termed *‘EctoShed’*, achieves expression of proteoforms of amine oxidase copper-containing 3 (AOC3) in adipocytes by capitalising on both the established lentiviral (LV) and AAV gene delivery systems, expressing full length or a soluble AOC3 mimic (m-sAOC3) isoforms. *In vitro* transduction of primary white adipocytes induced significant expression of both isoforms, retaining AOC3 enzymatic activity. *In vivo* delivery to inguinal white adipose tissue enabled depot-specific AOC3 expression and increased abundance of m-sAOC3 in the serum. Mice expressing m-sAOC3 exhibited reduced fat mass and fasting glucose levels.

**Conclusion:** *EctoShed* represents a versatile tool to dissect the functional roles of soluble proteoforms shed from adipocytes, enabling *in vitro* and *in vivo* applications in cardiometabolic health and disease.

## Background

Adipose tissue is an endocrine organ secreting proteins and peptides crucial for metabolic regulation (1–3). While adipokines such as leptin and adiponectin are released via the canonical ER-Golgi secretory pathway (4,5), others – including tumor necrosis factor alpha (TNF-α) and membrane proteins like amine oxidase copper-containing 3 (AOC3/vascular adhesion protein-1, VAP-1) – are secreted through a less studied mechanism, ectodomain shedding (6–8). Shedding thus controls the circulating levels and activity of surface proteins, and its dysregulation is linked to rheumatoid arthritis, Alzheimer’s disease, cancer and cardiometabolic disorders (8,9). Understanding the physiological role of shed proteoforms is hampered by the complexity of cleavage products and by tissue- and context-specific differences in cleavage kinetics. AOC3 is highly enriched in adipocytes, and both its transmembrane and soluble (sAOC3) isoforms exhibit amine oxidase and immunomodulatory activity (10). Elevated (s)AOC3 activity was reported in obesity, type 2 diabetes (T2D) and cardiovascular disease (CVD), highlighting its potential as a biomarker or therapeutic target (7,11,12). Notably, pharmacological inhibition of AOC3 in steatohepatitis (13) and in inflammatory conditions (14–16) has shown promising, but the functional contributions of individual AOC3 proteoforms remains unclear. Here, we present *EctoShed*, a novel *in vivo* combinatorial platform of adipocyte-resolved expression of transmembrane and soluble AOC3, offering broader utlity for studying shed proteins *in vivo*.

## Methods

### Animal care and research diets

All procedures were approved by the Danish Ministry of Environment and Agriculture (2023-15-0201-01544) and followed institutional guidelines. Male C57BL/6N mice (4 weeks old, Taconic Biosciences, Germany) were housed (3-4/cage) at 24□°C, 12-hour light/dark, with chow diet and water *ad libitum*. After 2 weeks of acclimatization, mice were switched to a low-fat diet (LFD; 10% kcal from fat, D12450J, Research Diets). Mice were then stratified by weight and randomly assigned to groups. At endpoint, mice were sacrificed by CO_2_ asphyxiation and cervical dislocation; Tissues were snap-frozen and stored at - 80 °C.

### Inguinal adipose tissue-directed AAV delivery

Recombinant AAV8 (1.5 × 10^12^ copies/mL PBS, VectorBuilder) was administered to 8-week-old male mice under isoflurane anesthesia (1-3%, 1L/min O_2_), as adapted from previous studies (17,18). Briefly, A <1 cm incision exposed posterior inguinal adipose tissue (iWAT), and each fat pad received 3-4 injections (total 40 μL, 6 x 10^10^ copies) using a 0.3 ml Micro Fine insulin syringe (BD). Analgesia (Rimadyl, 5 mg/kg) was provided daily for 72 h. Tissues were collected 6 weeks later.

### Body weight and glucose measurements

Body weight was recorded weekly. Fasting (6 h) glucose was measured from tail vein blood using a glucometer (Contour Next, Bayer).

### SVF isolation from inguinal white adipose tissue, adipogenesis and LV delivery

Stromal-vascular fraction (SVF) adipocyte precursor cells were isolated from iWAT from 4-6 weeks old male mice and differentiated into mature adipocytes as previously described (19). LV vectors (1.2 × 10□ cells/well, MOI 2.5, 10 µg/mL polybrene, VectorBuilder) were used to reverse transduce primary inguinal adipocytes (1° iWAd) at day 4 of differentiation and kept on FBS-free media from day 6 onwards. Transduction efficiency was confirmed by fluorescence after 24 h. Cells and culture media were harvested on day 8.

### Immunofluorescence staining and imaging

On day 4, LV-transduced 1° iWAd adipocytes were seeded onto sterilized 13 mm coverslips. On day 8 of differentiation, cells were fixed (acetone:methanol 1:1, -20 °C, 10 min) washed and blocked/permeabilized (PBS + 1% BSA + 0.1% Triton-X 100, RT for 30 min). Samples were incubated overnight (4 °C) with Alexa Fluor^TM^ 647-conjugated mouse anti-AOC3 (1:100, Santa Cruz, sc-166713), washed, nuclei stained with DAPI (5 ng/mL, 30 min, RT), mounted in Mowiol (Sigma-Aldrich, 81381), and imaged (Olympus BX53 Upright Fluorescence Microscope and CellSense software). Exposure settings were fixed. Negative controls (lacking primary antibody staining) were used to assess background noise. Mean fluorescence intensity was quantified in ImageJ (20) and normalized to DAPI and GFP controls.

### Cell culture media concentration

Cell culture media (supernatant) was collected on day 8 and concentrated using 50 kDa MWCO Amicon Ultra-2 Centrifugal FilterUnit (Millipore, UFC205024) at 4500 x g to obtain a volume of 100 – 300 μL.

### RNA isolation, cDNA synthesis and quantitative RT-PCR (qPCR)

RNA was isolated from adipocytes and tissues (iWAT, eWAT, BAT, liver), followed by cDNA synthesis and qPCR analysis as previously described (19). The following primers were used: Forward (FW): 5’-GTGGGATCTGAATGGAGAACTT-3’, Reverse (RV): 5’-CCTGTACCCTTGCCAATCAT-3’ (for *Gtf2b)*, FW: 5’-CTTCGGTGCTGGTGAGAAGT-3’, RV: 5’-TTCTTCAGCCCTGCCACATC-3’ (for *Aoc3)*; FW: 5’-GTTCTGCTGATTCTGCGAGC-3’, RV: 5’-AAGACAGAGGGGCAATGAAGA-3’ (for non-canonic sAOC3 mimic (*m-sAoc3). m-sAoc3* primers span the CDS for the adiponectin signal peptide (*SP*) and the proximal *Aoc3* exon to exclusively detect the soluble construct and not endogenous *Aoc3* or *Adiponectin*) FW: 5’-AAGAAGATGGTGCGCTCCTG-3’, RV: 5’-CTACCCCGACCACATGAAGC-3’ (for *Gfp)*.

Expression of *Gfp, Aoc3*, and m-*sAoc3* was normalized to Gtf2b using the 2^−ΔΔCt^ method (21).

### Quantification of AOC3 activity and protein abundance

Cell pellets and tissues were lysed in HES buffer (250 mM sucrose, 20 mM HEPES, 1 mM EDTA, with protease/phosphatase inhibitors (11873580001/4906845001 Roche). AOC3 activity was assayed in lysates and concentrated supernatants using the Amplex Red Monoamine Oxidase Assay Kit (Invitrogen, A12214) with monoamine oxidase A/B inhibition (clorgyline, pargyline) and subtraction of AOC3 residual activity in the presence of AOC3 inhibitor PXS-4681A (15). Fluorescence (RFU·min□^1^) was recorded over 30Lmin (CLARIOstar Plus plate reader). Values were normalized to protein content or sample volume. AOC3 protein content in serum and iWAT lysates was quantified by ELISA (Novus Biologicals, NBP2-78769) according to the manufacturer’s protocol and normalized to total protein or sample volume.

### Protein isolation, SDS-PAGE, and immunoblot analysis

iWAT was homogeneized and protein concentrations quantified as described elsewhere (19). 15 μ of protein was resolved on 7.5% SDS–PAGE gels (Bio-Rad) and transferred to nitrocellulose membranes (iBlot 2, Invitrogen). Membranes were blocked and probed with primary antibodies (anti-HSC70 1:2000 (sc-7298, Santa Cruz Biotechnology), anti-GFP 1:1000 (a11122, Invitrogen) and anti-AOC3 1:2000 (ab187202, Abcam) followed by washing and incubation with HRP-conjugated secondary antbibodies (anti-rabbit: #7074; anti-mouse: #7076; Cell Signaling Technology; 1:2000). Bands were visualized by ECL and quantified in ImageJ, normalized to loading controls.

### Statistical analysis

Data were analyzed using GraphPad Prism v10. Individual data points are shown. One-sample t-tests were used for comparison of fold-change differences and Kruskal-Wallis with Dunn’s post hoc for group comparisions. *p*□<□0.05 was considered significant.

## Results

We used LV (*in vitro*) and AAV serotype 8 (AAV8, *in vivo*) viral vectors to (over)express full-length AOC3 and its “mimic” soluble form sAOC3 (m-sAOC3), as well as green-fluorescent protein (GFP). Soluble AOC3 is believed to be generated by cleavage at the transmembrane anchor (22) and therefore the artificial m-sAOC3 construct was generated by eliminating the N-terminal amino acid sequence of the transmembrane isoform, thus lacking the intracellular (6 aa) and trans-membrane parts (21 aa) of the protein, while maintaining the proximal extracellular sequence of AOC3 that was fused to the Adiponectin secretion signal peptide (SP) (Fig. 1a, and Fig. 2a), thereby directing it to the ER lumen and secretion via the canonical signaling pathway into the extracellular milieu.

**Figure 1:**
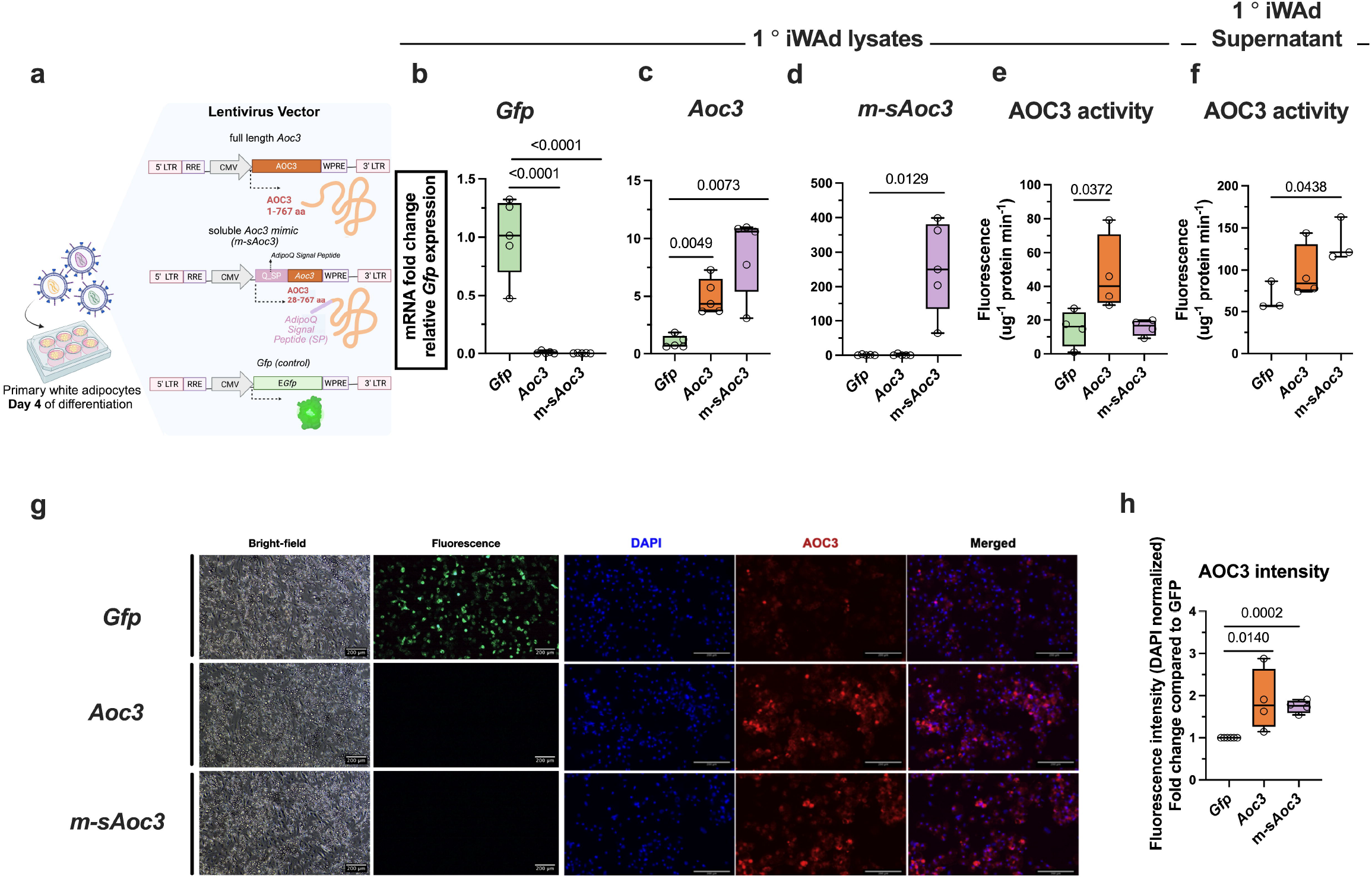
Lentiviral platform enables selective (over)expression of AOC3 proteoforms in primary adipocytes. (a) Representative scheme illustrating the LV constructs utilized for overexpression of AOC3 proteoforms upon transduction of SVF-isolated primary white adipocytes (1° iWad) on day 4 of differentiation: full length *Aoc3*; soluble *Aoc3 mimic* (m-*sAoc3) generated* by fusing Adiponectin signal peptide to the proximal extracellular sequence of *Aoc3*; and control construct expressing *Gfp*, under a CMV promoter. Generated using Biorender. Relative mRNA expression levels of (b) *Gfp*, (c) *Aoc3*, (d) m-*sAoc3* in 1° iWad lysates. Fluorometric determination of AOC3 enzyme activity in (e) 1° iWad lysates and (f) 1° iWad supernatant. (g) Representative bright-field images of 1° iWad transduced with LV vectors (left); representative immunofluorescent (IF) images of 1° iWad transduced with LV vectors followed by DAPI and AOC3 labelling and (h) quantification of AOC3 in 1° iWad. Quantification of IF shows individual points for independent experiments. Statistics: bar graphs represent the mean + s.e.m. with all data point represented. Unpaired, two-tailed Student’s test were performed (b,c,d), Non-parametric, Kruskal-Wallis with Dunn’s post hoc correction were performed (e,f,h).

**Figure 2:**
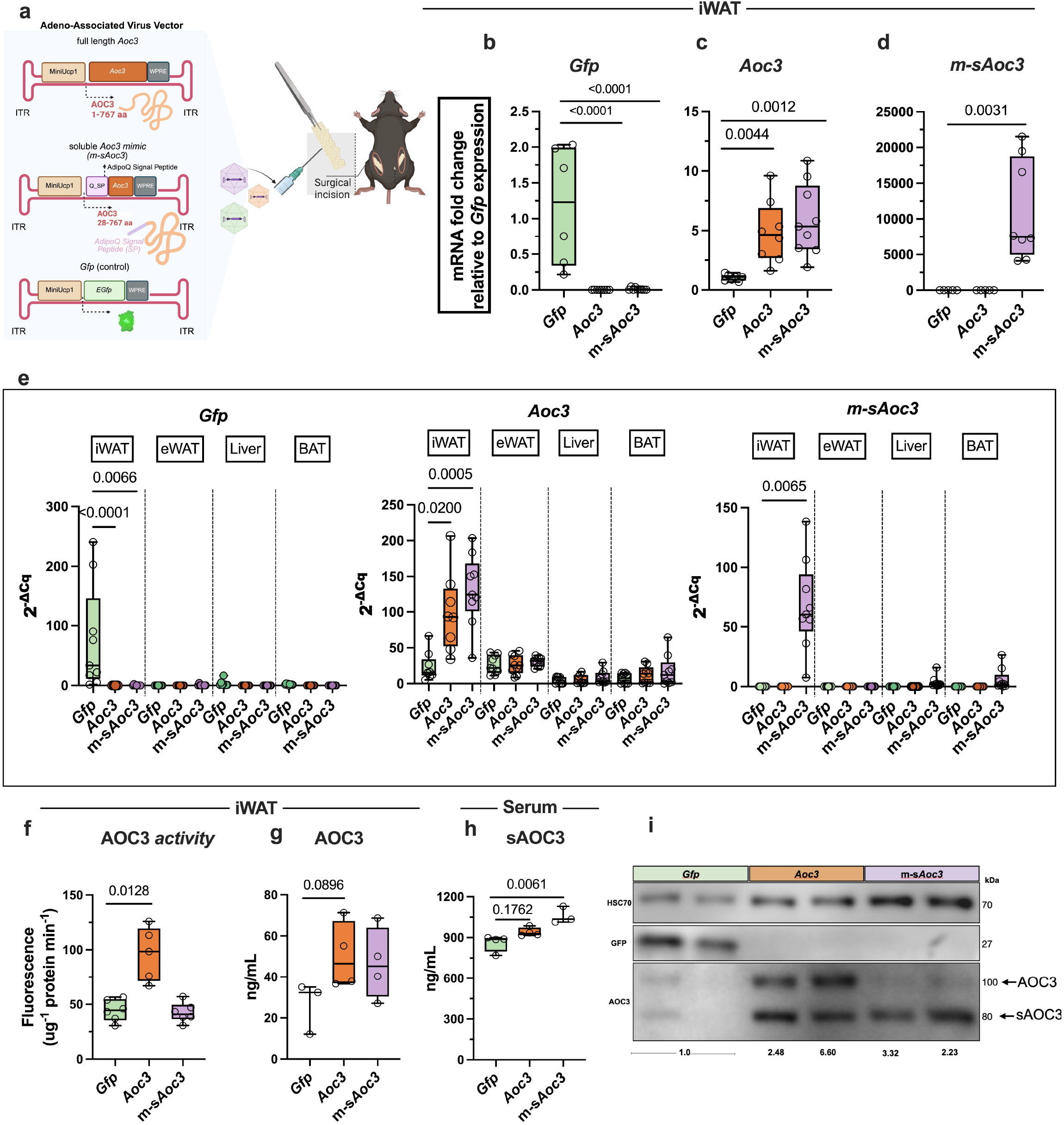
AAV platform enables iWAT-specific (over)expression and secretion into circulation of AOC3 proteoforms *in vivo*. Representative scheme illustrating the AAV constructs utilized for overexpression of AOC3 proteoforms in iWAT upon cirurgical preparation: full length *Aoc3*; soluble *Aoc3 mimic* (m-*sAoc3) generated* by fusing Adiponectin signal peptide to the proximal extracellular sequence of *Aoc3*; and control construct expressing *Gfp*, under a miniUcp1 promoter. Generated using Biorender. Relative mRNA expression levels of (b) *Gfp*, (c) *Aoc3, and* (d) m-*sAoc3* in iWAT. (e) Expression levels of indicated genes in iWAT, eWAT, liver and brown adipose tissue (BAT). (f) Fluorometric determination of AOC3 enzyme activity in iWAT and AOC3 protein quantification by enzyme-linked sorbent assay (ELISA) in (g) iWAT lysates and (h) serum. (i) Representative western blot showing GFP and AOC3 protein content in iWAT of transduced mice. HSC70 was used as loading control. Relative expression quantification is annotated in the figure. Statistics: bar graphs represent the mean + s.e.m. with all data point represented. Unpaired, two-tailed Student’s test were performed (b,c,d), Non-parametric, Kruskal-Wallis with Dunn’s post hoc correction were performed (e,f,g,h).

### LV transduction of primary inguinal adipocytes lead to robust *in vitro* overexpression of AOC3 proteoforms

LV-transduced 1° iWAd adipocytes (Fig. 1a) showed upregulation of *Gfp* (Fig. 1b), *Aoc3* (Fig. 1c) *and m-sAoc3* (Fig. 1d), with *Aoc3* expression increasing five to sevenfold while *m-sAoc3* mRNA exhibited a 200-fold increase. Given the dual role of AOC3 as an adhesion molecule and amine oxidase (10), retention of its enzyme activity is crucial for downstream applications of our platform. In accordance with the transcriptional over-expression, we observed increased AOC3 activity in the cell lysates following overexpression of *Aoc3* (Fig. 1e) and in the supernatant of the 1° iWAd expressing *m-sAoc3* (Fig. 1f), thus confirming functional protein secretion and enzyme activity. Fluorescence microscopy confirmed high transduction efficiency (as evidenced by GFP signal), and increased AOC3 protein abundance in cells (over)expressing *Aoc3* and m-*sAoc3* (Fig. 1g-h). Together, our results underscore the suitability of this platform for functional overexpression of shed-proteins in primary adipocytes.

### AAV-mediated over-expression of *m-sAoc3* in iWAT increases circulating m-sAOC3 in serum

We investigated the *in vivo* translational potential of this platform by testing the capacity for AAV-directed, iWAT-specific expression of m-sAOC3 in mice. AAV8 vectors encoding the same gene constructs, but under the control of synthetic *Ucp1* promoter (miniUcp1), enabling beige-adipose tissue specificity, were injected into iWAT, ensuring highly localized transduction (Fig. 2a). Expression analysis validated efficient induction of *Gfp* (Fig. 2b) and showed a five-fold increase of *Aoc3* (Fig. 2c) and m-*sAoc3* (Fig. 2d) expression compared to *Gfp* control. No appreciable increase in expression of both isoforms was detected in eWAT, BAT or liver tissues after 6 weeks of AAV administration, reflecting that the platform carries no risks of viral leakage into other metabolic tissues (Fig. 2e). In line with this, a two-fold increase in AOC3 enzyme activity in iWAT accompanied by increased protein levels following overexpression of *Aoc3*, but not *m-sAoc3*, compared to control was detected (Fig. 2f-g). Instead, expression of *m-sAoc3* elevated sAOC3 protein abundance in serum (Fig. 2h). Immunoblotting analysis confirmed i) increased AOC3 protein levels in iWAT from both constructs compared to the control group and ii) accumulation of a lower molecular weight form of AOC3 upon *m-sAOC3* expression *(Fig. 2i)*, in accordance to previously described sAOC3. (7).

### Increased levels of serum m-sAOC3 reduced body weight in mice

Increased circulating sAOC3 has been associated with obesity, atherosclerosis and CVD (11,23,24). We thus investigated the physiological impact of elevated sAOC3 mimic. 6 weeks after transduction, mice exhibited reduced body weight (**Fig. 3h**), likely secondary to reduced mass of iWAT (**Fig. 3i**) and eWAT (**Fig. 3j**). Aligned with this, animals showed lower fasting glucose (**Fig. 3k**). These results demonstrate, for the first time, that overexpression of sAOC3 mimic in iWAT, independent of its membrane-bound form, lowers body and fat mass and improves fasting glucose levels.

**Figure 3:**
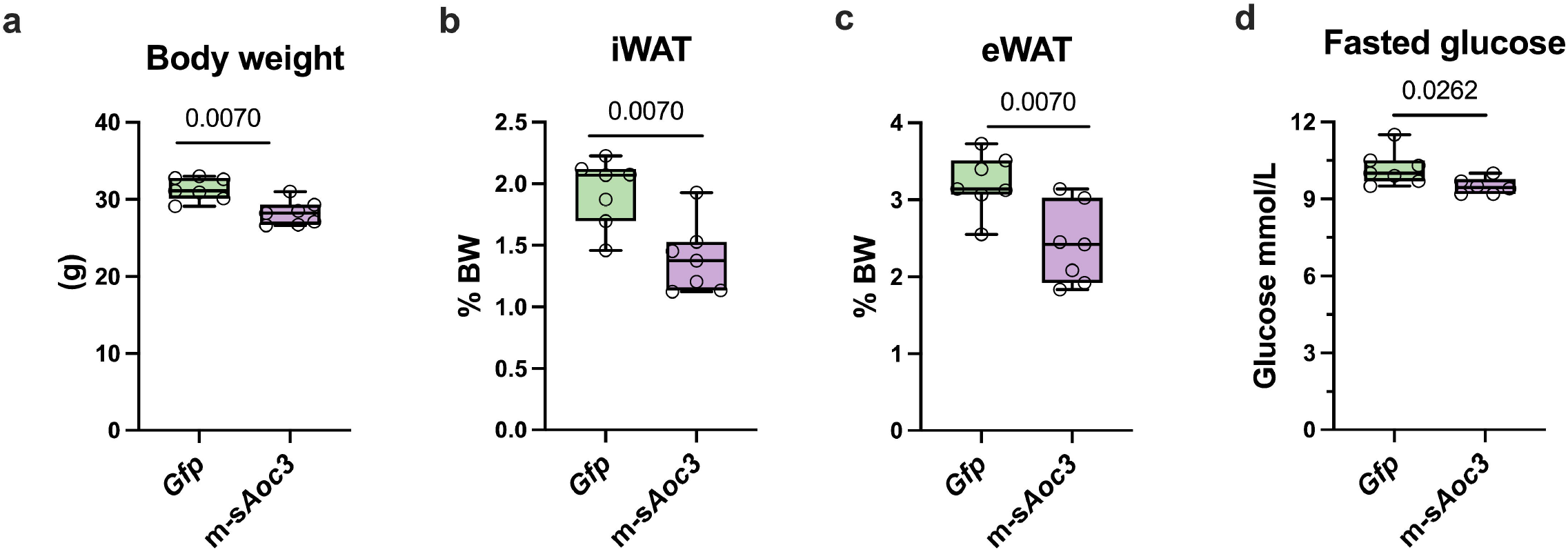
Overexpression of soluble AOC3 reduces body weight in mice. (a) Body weight (grams), (b) iWAT and (c) eWAT weight percentage relative to BW, and (d) fasted (6h) blood glucose levels 6 weeks after AAV-driven expression of m-s*Aoc3* versus *Gfp*. Statistics: bar graphs represent the mean + s.e.m. with all data point represented. Non-parametric, two-tailed Mann-Whitney t-test was performed (a,b,c,d).

## Discussion

Here, we describe a gene delivery platform enabling overexpression of transmembrane and their soluble proteoforms independently of endogenous protease (*sheddase*) activity, using AOC3 as a case point. Two separate gene-delivery platforms were employed based on the nature of the transduced adipocytes: LV vectors for primary adipocytes *in vitro* and AAV8 for transduction of adipocytes *in vivo*, leveraging their respective efficiencies and high transduction rates in adipocytes (25–29) to achieve depot-specific and robust over-expression of both AOC3 isoforms. While the use of adiponectin SP has been previously described in generating a glucagon-like peptide-1 analog fusion protein (30), this is the first application of this strategy for studying shed protein fragments. Importantly, by bypassing endogenous proteolytic regulation – which is influenced by post-translational modifications (8,9) - our system enables direct and consistent secretion of soluble proteoforms for mechanistic and physiological studies.

Prior *in vivo* models using (full length) AOC3 depletion or overexpression have been informative regarding AOC3 function, but limited by simultaneous presence of both isoforms (31–33). Our *EctoShed* platform circumvents this, providing selective expression of each proteoform and offering an alternative to traditional approaches like gain-of-function (GoF)/loss-of-function (LoF) or targeting metalloproteases. Notably, we show that increased sAOC3 mimic alone reduces body weight and fasting glucose, contrasting with reported positive correlation between tissue and serum sAOC3 levels and metabolic complications (7,11,34– 36). This may reflect context-dependent roles of sAOC3 in health and disease or that detrimental effects require the upregulation of both AOC3 proteoforms, opening new avenues to interrogate AOC3 function in metabolic regulation.

Altogether, our *EctoShed* platform facilitates the functional analysis of AOC3 proteoforms, and offers broader utility in the study of shed and secreted protein isoforms *in vivo*.

## List of abbreviations

(AAV): Adeno-associated viral
(AOC3): Oxidase copper-containing 3
(LV): Lentiviral
(VAP-1): Vascular adhesion protein-1
(TNF-α): Tumor necrosis factor alpha
(T2D): Type 2 diabetes
(CVD): Cardiovascular disease
(iWAT): Inguinal adipose tissue
(SVF): Stromal-vascular fraction
(1° iWAd): Primary inguinal adipocytes
(GoF): Gain-of-function
(LoF): Loss-of-function.

## Limitations

Limitations of this study: (1) The use of the miniUcp1 promoter enables specific over-expression in iWAT whereas it limits the use of systemic administration (i.p. injection) which could lead to transgene expression in brown adipose tissue, (2) also representing a limitation during thermoneutrality experiments. (3) The platform was not tested in obesity, where adipocyte hyperplasia could dilute AAV titers and require dose optimisation. (4) Our system allows expression of the proteforms above its already existing endogenous levels, requiring careful consideration when evaluating physiological effects due to possible suppraphysiological concentrations. One can thus speculate about the utility of this system during rescue experiments (such as GoF experiments in *Aoc3 knock-out* background), althought this was not tested in this study. (5) Many shed proteins remain poorly characterized, with unknown sheddases or cleavage sites (8,9). While this limits our ability to accurately mimic endogenous proteoforms, the system can be adapted to test alternative proteoform lenghts.

## Declarations

### Ethics approval

All animal procedures were carried out following institutional guidelines and were approved by the Danish Ministry of Environment and Agriculture (internal accession no. 2023-15-0201-01544).

### Availability of data and material

All data generated or analysed during this study are included in this published article. Information regarding the viral vectors and constructs used are available from the corresponding author on reasonable request.

## Acknowledgments

We thank Liv Holm, Stefanie Hansborg Kolstrup, Morten Ploug Kühlmann and Charlotte Laurfelt Munch Rasmussen for their assistance in animal experiments, Patricia S. S. Petersen and Brice Emanuelli for their support in training for surgical techniques.

## Consent for publication

Not applicable. This study does not involve human participants or individual person’s data.

## Author’s contributions

Conceptualization: HT, ART, NME, JWK were involved in the study and manuscript conceptualization and scientific discussion; Writing – original draft: ART, NME; Writing – review and editing, corrections and comments: ART, NME, HT and JWK. Experimental procedures performed by ART, NME, NS.

## References

1. Kershaw EE, Flier JS. Adipose tissue as an endocrine organ. In: Journal of Clinical Endocrinology and Metabolism. 2004. p. 2548–56.

2. Cypess AM. Reassessing Human Adipose Tissue. New England Journal of Medicine. 2022 Feb 24;386(8):768–79.

3. Pogodziński D, Ostrowska L, Smarkusz-Zarzecka J, Zyśk B. Secretome of Adipose Tissue as the Key to Understanding the Endocrine Function of Adipose Tissue. Vol. 23, International Journal of Molecular Sciences. MDPI; 2022.

4. Xie L, Boyle D, Sanford D, Scherer PE, Pessin JE, Mora S. Intracellular trafficking and secretion of adiponectin is dependent on GGA-coated vesicles. Journal of Biological Chemistry. 2006 Mar 17;281(11):7253–9.

5. Xie L, O’Reilly CP, Chapes SK, Mora S. Adiponectin and leptin are secreted through distinct trafficking pathways in adipocytes. Biochim Biophys Acta Mol Basis Dis. 2008 Feb;1782(2):99–108.

6. Tien WS, Chen JH, Wu KP. SheddomeDB: The ectodomain shedding database for membrane-bound shed markers. BMC Bioinformatics. 2017 Mar 14;18.

7. Abella A, García-Vicente S, Viguerie N, Ros-Baró A, Camps M, Palacín M, et al. Adipocytes release a soluble form of VAP-1/SSAO by a metalloprotease-dependent process and in a regulated manner. Diabetologia. 2004 Mar;47(3):429–38.

8. Lichtenthaler SF, Lemberg MK, Fluhrer R. Proteolytic ectodomain shedding of membrane proteins in mammals—hardware, concepts, and recent developments. EMBO J. 2018 Aug;37(15).

9. Lichtenthaler SF, Meinl E. To cut or not to cut: New rules for proteolytic shedding of membrane proteins. Journal of Biological Chemistry. 2020 Aug 28;295(35):12353–4.

10. Marko Salmi* Sirpa Jalkanen. VAP-1: an adhesin and an enzyme. Trends Immunol. 2001;22(4):211–6.

11. Yang H, Liu CN, Wolf RM, Ralle M, Dev S, Pierson H, et al. Obesity is associated with copper elevation in serum and tissues. Metallomics. 2019 Aug 1;11(8):1363–71.

12. Fasolato S, Bonaiuto E, Rossetto M, Vanzani P, Ceccato F, Vittadello F, et al. Serum Vascular Adhesion Protein-1 and Endothelial Dysfunction in Hepatic Cirrhosis: Searching for New Prognostic Markers. Int J Mol Sci. 2024 Jul 1;25(13).

13. Newsome PN, Sanyal AJ, Neff G, Schattenberg JM, Ratziu V, Ertle J, et al. A randomised Phase IIa trial of amine oxidase copper-containing 3 (AOC3) inhibitor BI 1467335 in adults with non-alcoholic steatohepatitis. Nat Commun. 2023 Dec 1;14(1).

14. Jarnicki AG, Schilter H, Liu G, Wheeldon K, Essilfie AT, Foot JS, et al. The inhibitor of semicarbazide-sensitive amine oxidase, PXS-4728A, ameliorates key features of chronic obstructive pulmonary disease in a mouse model. Br J Pharmacol. 2016;173(22):3161–75.

15. Foot JS, Yow TT, Schilter H, Buson A, Deodhar M, Findlay AD, et al. PXS-4681A, a potent and selective mechanism-based inhibitor of SSAO/VAP-1 with anti-inflammatory effects in vivo. Journal of Pharmacology and Experimental Therapeutics. 2013 Nov;347(2):365–74.

16. Schilter HC, Collison A, Russo RC, Foot JS, Yow TT, Vieira AT, et al. Effects of an anti-inflammatory VAP-1/SSAO inhibitor, PXS-4728A, on pulmonary neutrophil migration. Respir Res. 2015 Mar 20;16(1).

17. Sveidahl Johansen O, Ma T, Hansen JB, Markussen LK, Schreiber R, Reverte-Salisa L, et al. Lipolysis drives expression of the constitutively active receptor GPR3 to induce adipose thermogenesis. Cell. 2021 Jun 24;184(13):3502–3518.e33.

18. Modica S, Straub LG, Balaz M, Sun W, Wolfrum C, Varga L, et al. Bmp4 Promotes a Brown to White-like Adipocyte Shift. Cell Rep. 2016 Aug 23;16(8):2243–58.

19. Khani S, Topel H, Kardinal R, Tavanez AR, Josephrajan A, Larsen BDM, et al. Cold-induced expression of a truncated adenylyl cyclase 3 acts as rheostat to brown fat function. Nat Metab. 2024 Jun 1;6(6):1053–75.

20. Rueden CT, Schindelin J, Hiner MC, DeZonia BE, Walter AE, Arena ET, et al. ImageJ2: ImageJ for the next generation of scientific image data. BMC Bioinformatics. 2017 Nov 29;18(1).

21. Livak KJ, Schmittgen TD. Analysis of relative gene expression data using real-time quantitative PCR and the 2-ΔΔCT method. Methods. 2001;25(4):402–8.

22. Murata M, Noda K, Kawasaki A, Yoshida S, Dong Y, Saito M, et al. Soluble Vascular Adhesion Protein-1 Mediates Spermine Oxidation as Semicarbazide-Sensitive Amine Oxidase: Possible Role in Proliferative Diabetic Retinopathy. Curr Eye Res [Internet]. 2017 Dec 2 [cited 2023 Nov 25];42(12):1674–83. Available from: https://www.tandfonline.com/doi/abs/10.1080/02713683.2017.1359847

23. Li H, Du S, Niu P, Gu X, Wang J, Zhao Y. Vascular Adhesion Protein-1 (VAP-1)/Semicarbazide-Sensitive Amine Oxidase (SSAO): A Potential Therapeutic Target for Atherosclerotic Cardiovascular Diseases. Vol. 12, Frontiers in Pharmacology. Frontiers Media S.A.; 2021.

24. Papukashvili D, Rcheulishvili N, Deng Y. Attenuation of weight gain and prevention of associated pathologies by inhibiting SSAO. Vol. 12, Nutrients. MDPI AG; 2020.

25. Behrens J, Braren I, Jaeckstein MY, Lilie L, Heine M, Sass F, et al. An efficient AAV vector system of Rec2 serotype for intravenous injection to study metabolism in brown adipocytes in vivo. Mol Metab. 2024 Oct 1;88.

26. Merentie M, Lottonen-Raikaslehto L, Parviainen V, Huusko J, Pikkarainen S, Mendel M, et al. Efficacy and safety of myocardial gene transfer of adenovirus, adeno-associated virus and lentivirus vectors in the mouse heart. Gene Ther. 2016 Mar 1;23(3):296–305.

27. O’neill SM, Hinkle C, Chen SJ, Sandhu A, Hovhannisyan R, Stephan S, et al. Targeting adipose tissue via systemic gene therapy. Gene Ther. 2014;21(7):653–61.

28. Bates R, Huang W, Cao L. Adipose Tissue: An Emerging Target for Adeno-associated Viral Vectors. Vol. 19, Molecular Therapy Methods and Clinical Development. Cell Press; 2020. p. 236–49.

29. Boychenko S, Abdullina A, Laktyushkin VS, Brovin A, Egorov AD. Assessment of Adipocyte Transduction Using Different AAV Capsid Variants. Pharmaceuticals. 2024 Sep 1;17(9).

30. Zhao T, Lv J, Zhao J, Huang X, Xiao H. Expression of Human Globular Adiponectin-Glucagon-Like Peptide-1 Analog Fusion Protein and Its Assay of Glucose-Lowering Effect In Vivo [Internet]. Vol. 8, Int. J. Med. Sci. 2011. Available from: http://www.medsci.org203

31. Bour S, Caspar-Bauguil S, Iffiú-Soltész Z, Nibbelink M, Cousin B, Miiluniemi M, et al. Semicarbazide-sensitive amine oxidase/vascular adhesion protein-1 deficiency reduces leukocyte infiltration into adipose tissue and favors fat deposition. American Journal of Pathology. 2009;174(3):1075–83.

32. Grès S, Bour S, Valet P, Carpéné C. Benzylamine antihyperglycemic effect is abolished by AOC3 gene invalidation in mice but not rescued by semicarbazidesensitive amine oxidase expression under the control of aP2 promoter. J Physiol Biochem. 2012 Dec;68(4):651–62.

33. Jargaud V, Bour S, Tercé F, Collet X, Valet P, Bouloumié A, et al. Obesity of mice lacking VAP-1/SSAO by Aoc3 gene deletion is reproduced in mice expressing a mutated vascular adhesion protein-1 (VAP-1) devoid of amine oxidase activity. 2021; Available from: www.genoway.com;

34. Aalto K, Maksimow M, Juonala M, Viikari J, Jula A, Kähönen M, et al. Soluble vascular adhesion protein-1 correlates with cardiovascular risk factors and early atherosclerotic manifestations. Arterioscler Thromb Vasc Biol. 2012 Feb;32(2):523–32.

35. Kuo CH, Wei JN, Yang CY, Ou HY, Wu HT, Fan KC, et al. Serum vascular adhesion protein-1 is up-regulated in hyperglycemia and is associated with incident diabetes negatively. Int J Obes. 2019 Mar 1;43(3):512–22.

36. Kurkijarvi R, Yegutkin GG, Gunson BK, Jalkanen S, Salmi M, Adams DH. Circulating soluble vascular adhesion protein 1 accounts for the increased serum monoamine oxidase activity chronic liver disease. Gastroenterology. 2000;119(4):1096–103.

